# *TraceMontage*: a Method for Merging Multiple Independent Neuronal Traces

**DOI:** 10.1101/703900

**Authors:** Aslan S Dizaji, Logan A Walker, Dawen Cai

## Abstract

**Background:** The ability to reconstruct neuronal networks, local microcircuits, or the entire connectome is a central goal of modern neuroscience. Recently, advancements in sample preparation (e.g., sample expansion and Brainbow labeling) and optical (e.g., confocal and light sheet) techniques have enabled the imaging of increasingly large neural systems with high contrast. Tracing neuronal structures from these images proves challenging, however, necessitating tools that integrate multiple neuronal traces, potentially derived by various methods, into one combined (*montaged*) result.

**New Method:** Here, we present TraceMontage, an ImageJ/Fiji plugin for the combination of multiple neuron traces of a single image, either redundantly or non-redundantly. Internally, it uses graph theory to connect topological patterns in the 3-D spatial coordinates of neuronal trees. The software generates a single output tracing file containing the montage traces of the input tracing files and provides several measures of consistency analysis among multiple tracers.

**Results and Comparison to existing method(s):** To our knowledge, our software is the first dedicated method for the combination of tracing results. Combining multiple tracers increases the accuracy and speed of tracing of densely-labeled samples by harnessing collaborative effort. This utility is demonstrated using fluorescence microscope images from the hippocampus and primary visual cortex (V1) in Brainbow-labeled mice.

**Conclusions:** *TraceMontage* provides researchers the ability to combine neuronal tracing data generated by either the same or different method(s). As datasets become larger, the ability to trace images in this parallel manner will help connectomics scale to increasingly larger neural systems.

## 1. Introduction

One of the major goals of neuroscience is to map neuronal circuits at the microscopic resolution to elucidate their structure-function relationship and elaborate on the connection of brain and behavior, also known as connectomics. For decades, these studies have been conducted through direct observation of neuronal morphology under light and electron microscopy.

Traditionally, neurons are sparsely labeled to be unambiguously captured in a single sample. It leads directly to the need for computational tools to identify and separate neural structures from image stacks. In 2010, the *DIADEM* (DIgital reconstruction of Axonal and DEndritic Morphology) contest was organized to encourage development in the study of 3D reconstructions of neurons. Recently, the *BigNeuron* (Peng et al., 2015) project was formed, aiming to perform comprehensive comparisons of more recent developments in the field using standardized data protocols and evaluation methods. These two projects have resulted in a burgeoning of automated methods for producing neuronal reconstructions. Additionally, manual and semi-manual approaches have been created which allow human annotation of images to form 3D reconstructions. These methods include *Simple Neurite Tracer* (Longair et al., 2011), *Neuromantic (Myatt et al., 2012)* or commercial tools such as *Neurolucida* and *iMaris*. More recently, the transgenic Brainbow techniques (Cai et al., 2013; Lichtman et al., 2008; Livet et al., 2007) have facilitated high throughput neuronal imaging at the microscale resolution, using the combinatorial expression of several fluorescent proteins in neurons to color-tag individual cells and fluorescence microscopy to visualize them. This allows dense and unique labeling of neuronal processes, which permits short-to-long-range circuit tracing of multiple neurons of the same sample by the software *nTracer* (Roossien et al., 2019).

The digitization of neuronal morphology normally describes the tree-like branching of axons and dendrites as a sequence of interconnected cylinders (as in the industry-standard SWC file format). In this sparse representation, each point in the arbor is usually characterized by five values including the three Euclidean coordinates, the diameter of the cylinder, and the identity of the “parent” point from which it originates. While the labor-intensive and rate-determining process of neuronal reconstruction is largely facilitated by increasingly automated computational algorithms, error checking and quality control still require human intervention (Parekh and Ascoli, 2013). In general, automated solutions to neuronal reconstructions aim to answer the three interrelated areas of active study (Acciai et al., 2016): the morphological characterization of cell types, mapping projections of single cells in the whole brain, and describing convergence-divergences patterns in the neuronal networks of different brain regions. The first and second problems demand developing image processing techniques to automate neuron tracing in single-color samples of brain tissues. Brainbow, as a potential candidate to investigate the third problem, intends to map the convergence-divergences patterns in neuronal networks, e.g. among excitatory pyramidal neurons and different types of inhibitory interneurons.

Despite the increasing amount of data produced through these methods, utilities for the postprocessing of SWC files have been notably lacking. A common operation, the merging of tracing results, is often done manually in annotation software, requiring the effort of skilled technicians. The requirement that tracing between multiple images be merged can arise from several situations, for example, if multiple (overlapping) tiles of a wide-field image are traced independently (e.g. to increase throughput), if different neurons of a single image are traced independently, or if a single neuron in a single image is traced redundantly as a quality control measure. To address these different scenarios, we present *TraceMontage*, an ImageJ/Fiji plugin, which was developed to provide an automated combination (montage) methods for merging the neural traces generated by multiple tracers or modalities. These tracing datasets belong to one image or two adjacent images. In the former case, the plugin is also able to remove the redundant traces of one tracing dataset, and, in the latter case, the two adjacent images must have an overlapped region. *TraceMontage* uses graph theory to find patterns in the 3-D spatial coordinates of neuronal trees and uses a model with few free parameters. The algorithms applied are fast and capable of handling large tracing datasets provided in SWC format (which contain the neurites’ x, y, and z coordinates with a known convention for branch naming in the form of a full binary tree structure). It is applicable to both monochromatic neuron labeling and multi-spectral neuron labeling (e.g., Brainbow) and forms a key part of the growing toolkit of connectomics data analysis. In the following sections, we will demonstrate *TraceMontage* using results generated by *nTracer* from multi-spectral Brainbow samples. The Brainbow images provide color information for the identification of distinct neurons, which serves as a more general example than the canonical monochrome images.

## 2. Methods

### 2.1. The TraceMontage Workflow

Figure 1 gives an overview of the workflow automated by *TraceMontage*. To obtain the color of branches, *TraceMontage* accepts one or two TIFF-formatted image files, while tracings are read from one or two files in SWC format or *nTracer-ZIP* format. For each branch in each image, overlapping traces between the input files are identified using their spatial coordinates and the color of the traces (if the TIFF file provided has multiple channels of labeling, such as an ImageJ hyperstack). Next, branches which overlap are merged locally to remove redundant data, and, finally, the trees which contain the overlapping branches are combined into a single model, preserving the binary structure initiated from a single soma node. Notably, in cases of disagreement between two tracers, the software will default to the “Primary” tracer, which is indicated by the first input tracing file; this can result in differences of merging results based on the selection of the primary. After running, the plugin generates one output tracing data file which contains the montage traces of the input tracing file(s). Additionally, a range of quality control metrics that describe the consistency between tracers is calculated and presented upon completion.

**Figure 1:**
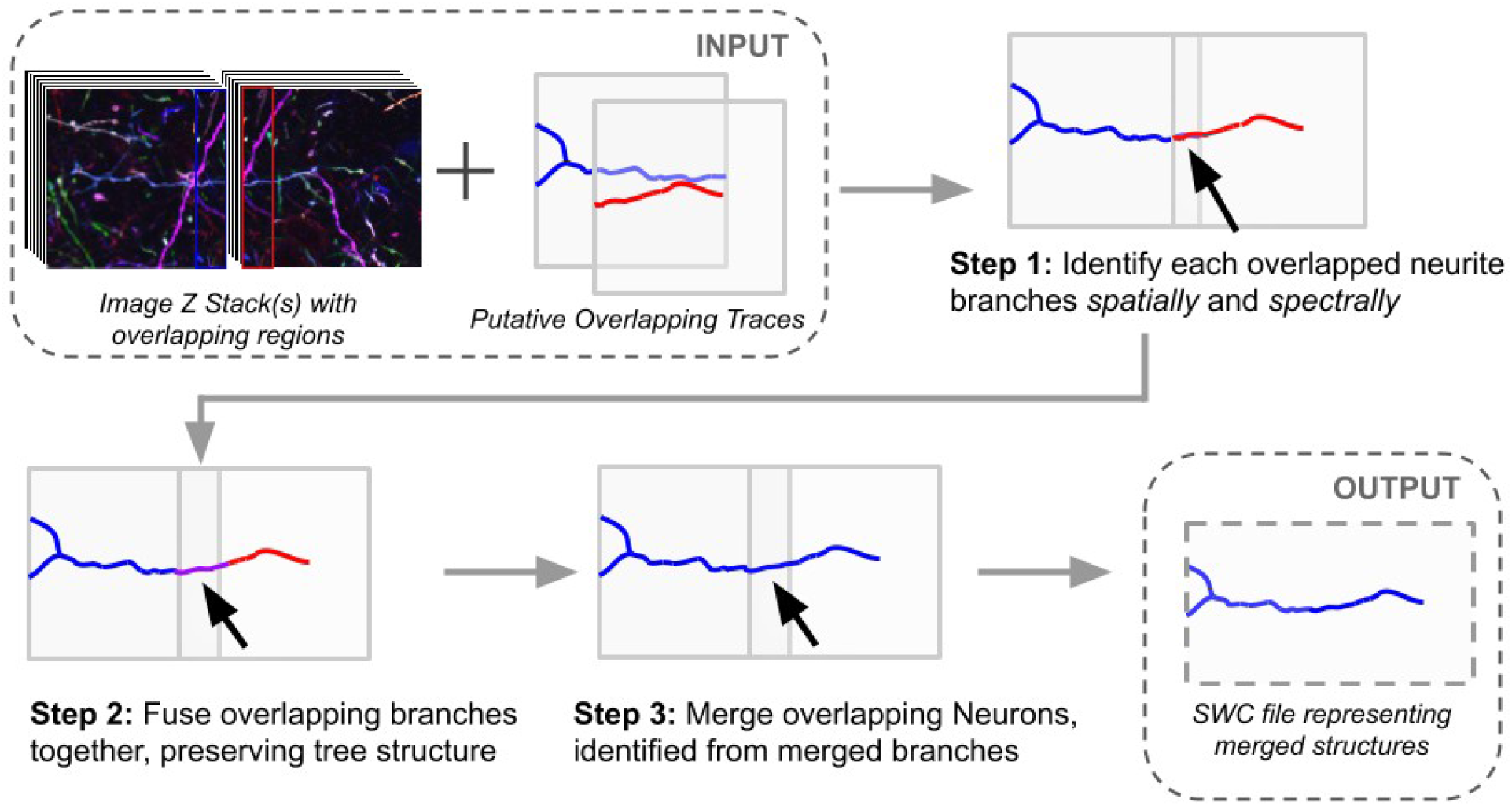
An overview of the TraceMontage workflow. First, the user inputs two overlapping neuron traces (generated with either automatic or manual methods) and corresponding microscope image stacks. The software identifies neuron traces which are overlapping via graph theory methods, overlapping branches are merged together, and, finally, neurons which contain overlapping branches are merged. The merged result is output for visualization or additional analysis.

### 2.2. Algorithms

*TraceMontage* utilizes a theoretical model for neural reconstruction derived on that used in the *nTracer* software (**Figure S1A**). This model represents a neuron reconstruction as a tree graph, which has a root node, leaf nodes, branching nodes, and other internodes, following the definitions of (Peng et al., 2011). This allows the manipulation of these structures within the framework of graph theory. As such, *TraceMontage* implements a fuzzy-logic (Radojevic and Meijering, 2017) inspired algorithm for the identification of overlapping branches. Additional, detailed information about the algorithms implemented in *TraceMontage* can be found in the **Supplementary Methods**.

### 2.3. Testing and Demonstration

*TraceMontage* was tested under various scenarios to demonstrate its utility. Test microscope images were collected from Brainbow-labeled mouse Hippocampus or primary visual cortex (V1) samples, as indicated in (Roossien et al., 2019; Roossien and Cai, 2017), and split into overlapping tiles to simulate the result of a two-exposure imaging experiment. Each tile was independently traced with *nTracer* by separate, trained biologists (see **Acknowledgements**), followed by merging with *TraceMontage*. Resulting traces were then annotated for correctness through comparisons to a “gold-standard” traced from the original (before splitting) image.

### 2.4 Software Availability

The *TraceMontage* plugin, example Brainbow images and tracing results are available from the Cai Lab website at https://www.cai-lab.org/tracemontage. *TraceMontage* is under GPL3 License (https://www.gnu.org/licenses/gpl-3.0.en.html). After installation, the software can be started from ImageJ/Fiji, under the “plugins” interface.

## 3. Results

### 3.1. Merging Horizontally-aligned Image Tiles

In **Figure 2**, image tiles of Parvalbumin-expressing Basket Cells (PVBCs) in the mouse Hippocampus were traced, followed by merging with *TraceMontage*. As demonstrated in the enlarged portion of the figure, *TraceMontage* is capable of merging matching traces in the overlapped region of two Brainbow images. It is notable, however, that inaccuracies in the human tracing (possibly caused by the difficulty imposed by the dense labeling) can lead to an incomplete merging of all possible neurites. The default values of parameters were used in this example, although optimization of the free parameters could lead to decreases in the number of merged traces, at the cost of stringency and/or accuracy of the combined set of output neurons.

**Figure 2:**
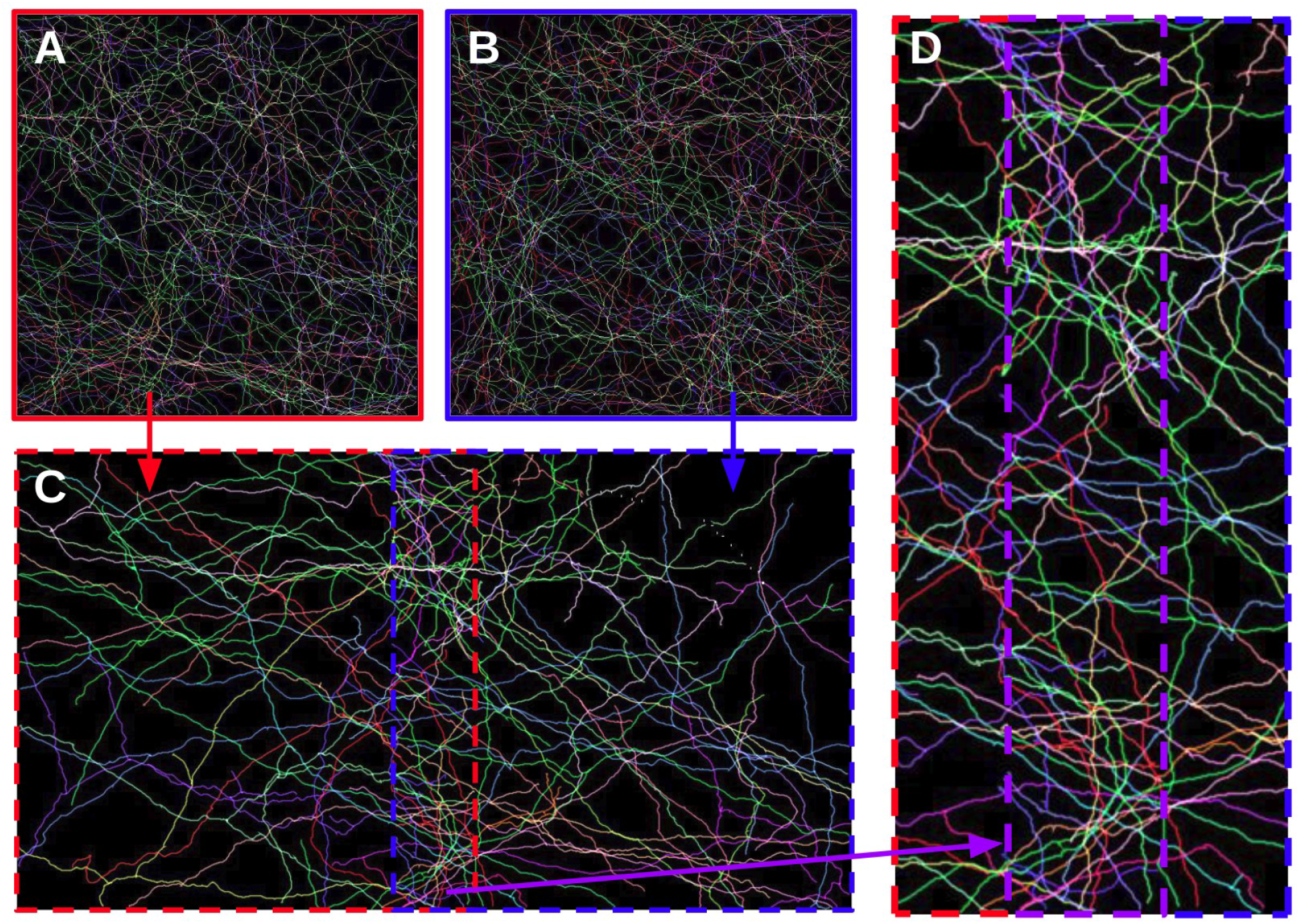
An example merge performed with TraceMontage. An image of Parvalbumin-expressing basket cells (fast-spiking inhibitory interneurons) of the Hippocampus of mouse brain was split into two overlapping tiles; (**A and B**) All visible neurites in each tile was semi-manually traced using the *nTracer* tool by two different scientists; **(C)** The traces with overlapping region were shown after merging using *TraceMontage*; **(D)** The overlapping and adjacent regions are enlarged to show the merged traces clearly connect between (A) and (B). All panels are Z-projected renderings of the path, colored to match the original sample’s Brainbow labeling; the image is 560 x 560 x 225 pixels total.

### 3.2 Additional Examples

*TraceMontage* was tested using image tiles from a vertically-aligned PVBC image (**Figure S6**), multiple tracers on a single PVBC image (**Figure S7 and S8**). These cases were chosen to demonstrate the use of *TraceMontage* as a tool for tracing wide-field images (similar to **Figure 1**), and as an automated proofreading tool between multiple tracers, respectively.

## 4. Discussion

Increasing computational power and advancements in imaging technology continue to enable the collection and processing of large connectomic images. Currently, one rate-determining step in the usage of this data is to efficiently and consistently generate tracing results from the whole volume of very large microscopic images. The algorithms powering *TraceMontage* are able to provide an efficient solution for the combination of tracing results, and, as a result, this plugin and its algorithms make a contribution to the large-scale neuronal tracing in computational connectomics.

There are, however, some limitations regarding the use of this software. First, in the current format of its algorithms, the plugin is only applicable to the alignment of neurites representable with a full binary tree structure. This prohibits its usage to other non-binary structures, such as the vasculature system (CITE some of those tracing/segmentation results). This issue can be resolved by considering the topological relations between the number of nodes and edges when the tree structure is not necessarily binary. A second limitation is that, in some situations, some traces may not be combined. This is most evident when there are neurites that were not traced in both input reconstructions. In that case, some original tracings may be excluded (for consistency) from the final montage result, and therefore, some information which one of the tracers has been provided would be missed. This issue arises because the two end-points of each branch are the only signatures of the branch and there is no referencing to the internodes of the branch in the whole tree. We think this could be addressed in a future version of the software, but would be computationally more expensive due to the larger number of comparisons which would be required. A third limitation is that the current version arbitrarily assumes one tracing result is always correct when presented differences. The future version may provide options to calculate an average as the merged results. Finally, new algorithms for merging more than two tracing results would be powerful for generating “gold-standard” with higher accuracy in a large collaborative tracing setting.

## Author Contributions

DC conceived the project. AD developed the algorithms and the plugin. AD, DC, and other members of the Cai Lab provided the neuronal tracing data. AD wrote the first draft of the manuscript. All authors contributed to manuscript revision, read, and approved the final draft.

## Acknowledgements

We would like to thank Douglas Roossien and Benjamin Sadis of the Cai Lab for providing the neuronal tracing data and test images. We would like to thank Hamid Soltanian-Zadeh and other members of the SCS-IPM (School of Cognitive Sciences of the Institute for Research in Fundamental Sciences in Tehran, Iran) for the help during the course of the project and thank Alireza Rahmat-Abadi for the help with visualization.

## Supplementary Material

This article has accompanying supplementary material.

