## Supplemental Material for "*TraceMontage*: a Method for Merging Multiple Independent Neuronal Traces"

### Supplemental Figures

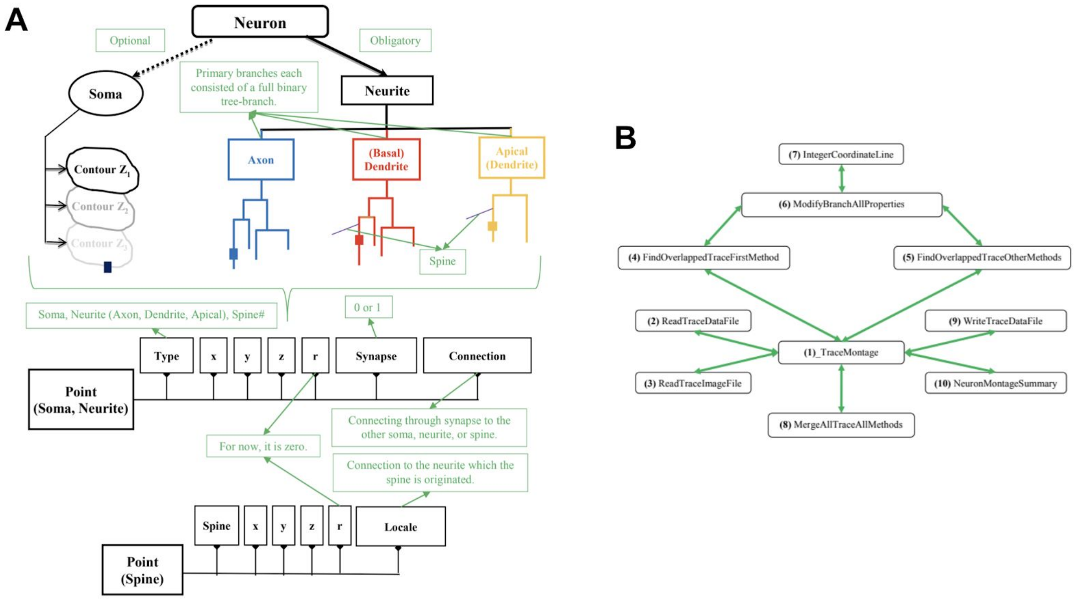


**Figure S1. Schematics of *TraceMontage* data structure and program diagram.**
**(A)** An overview of *TraceMontage* data model is shown. It is composed of soma contours, neurites with a full binary tree structure (axons, basal dendrites, apical dendrites), spines, and synapses. This structural format is the same as the structural format of the output of *nTracer (Roossien et al., 2019)*, another ImageJ/Fiji plugin, which was particularly developed for the neuron tracing of Brainbow images; **(B)** A schematic diagram showing the interactions among ten classes of the *TraceMontage* program.

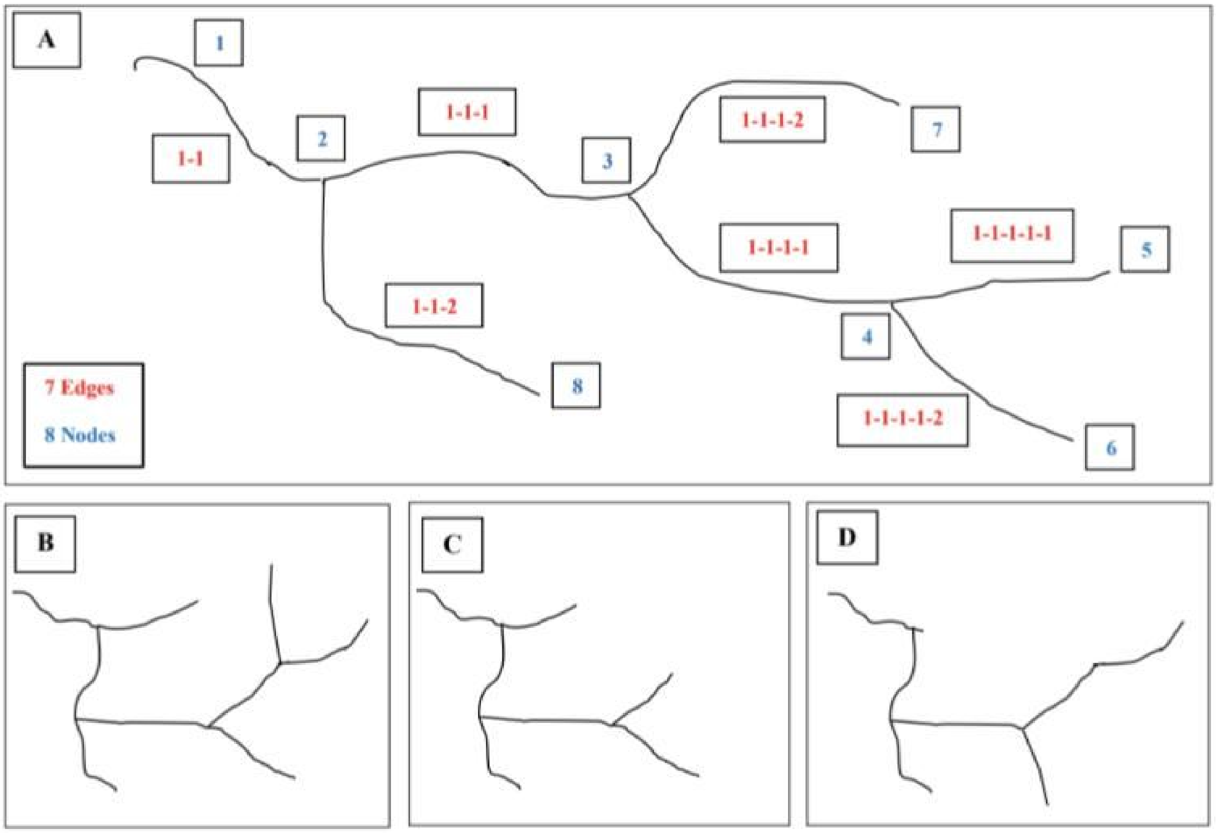


**Figure S2. Graphic model of the neuron tracing result as a binary tree.**
**(A)** The relations between all edges and nodes of a tree branch. The graphical definitions of "accurate" and "complete" traces for a tree: **(B)** "accurate" and "complete"; **(C)** "accurate" and "incomplete"; **(D)** "inaccurate" and "incomplete".

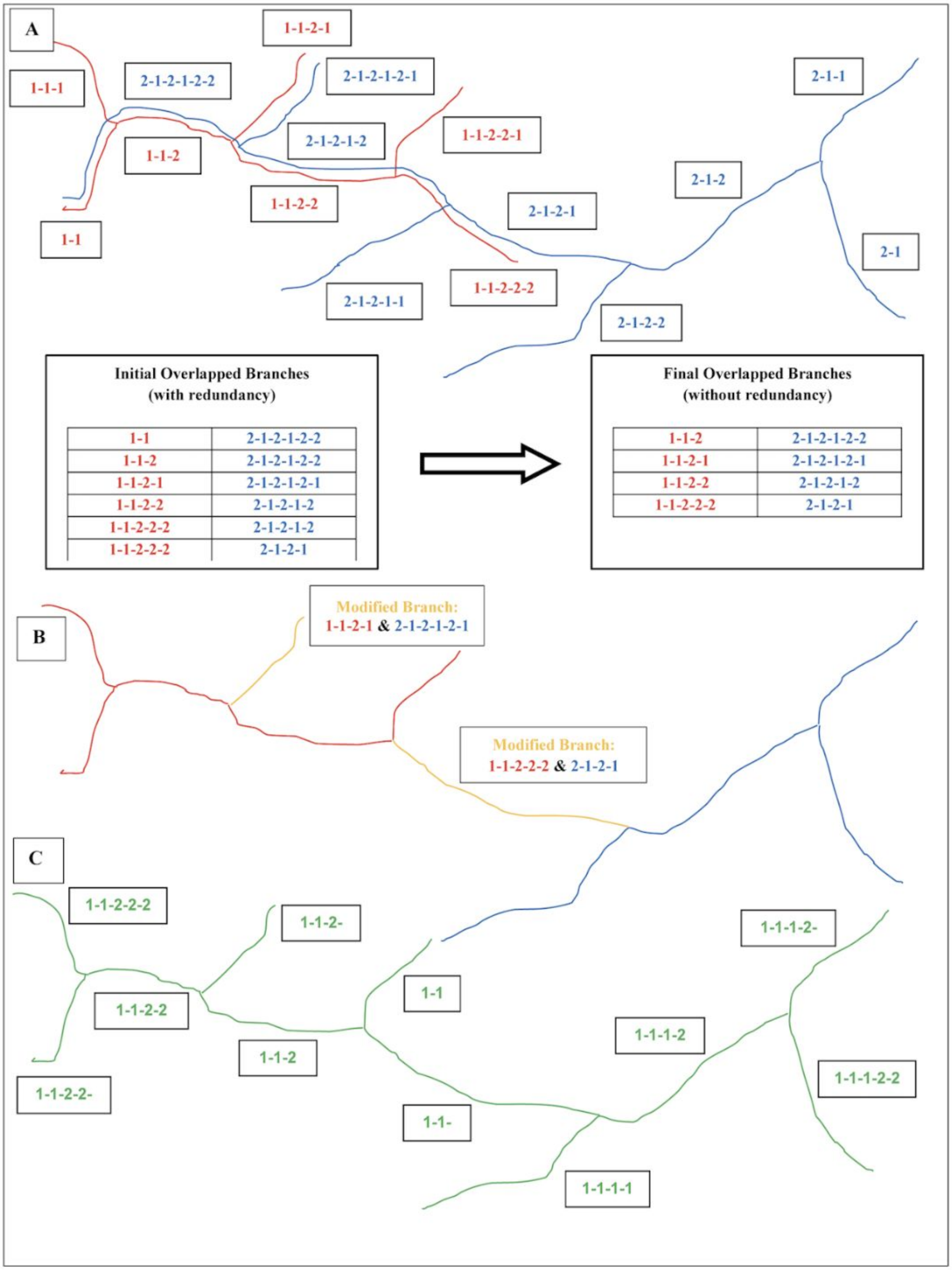


**Figure S3. A schematic example of the steps for merging two overlapping branches.**
**(A)** Identify the overlapped branches of two overlapped tree branches. **(B)** Modify the coordinates of the overlapped branches. **(C)** Merge the two overlapping trees to create the final montaged tree.

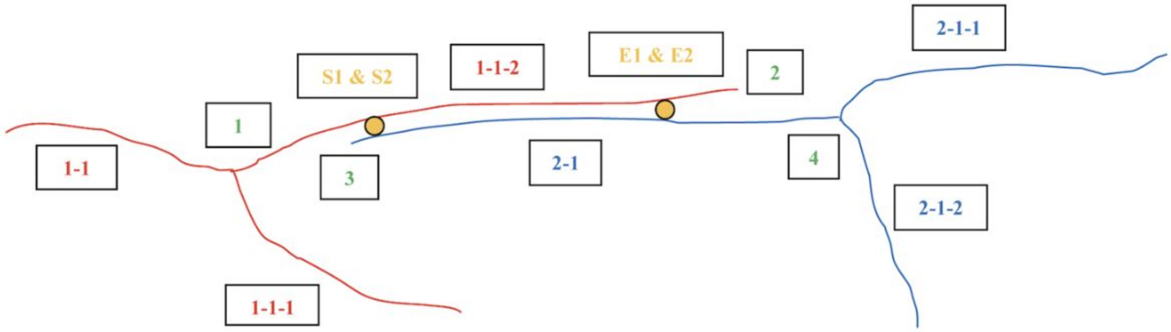


**Four Topological Variables:**
**(1-1-2 & 2-1)**

flag1: 1
flag2: 0
flag3: 0
flag4: 2

**Four Indices:**
**(1-1-2 & 2-1)**

S1: indexStart1
S2: indexStart2
E1: indexEnd1
E2: indexEnd2

**1 & 4 are the two selected end points of the two overlapped branches for preservation.**

**The Overlapped Branch 1-1-2:**

flag1: 1 & flag2: 0

| | Index | X | Y | Z |
|---|---|---|---|---|
| 1 | 0 | 23 | 43 | 12 |
| | 1 | 24 | 43 | 12 |
| | … | … | … | … |
| S1 | 54 | 63 | 50 | 19 |
| | 55 | 64 | 50 | 20 |
| | … | … | … | … |
| E1 | 123 | 98 | 66 | 26 |
| | 124 | 99 | 67 | 26 |
| | … | … | … | … |
| 2 | 161 | 143 | 87 | 32 |
| | 162 | 144 | 88 | 33 |

S1 < E1 (always)

**The Overlapped Branch 2-1:**

flag3: 0 & flag4: 2

| | Index | X | Y | Z |
|---|---|---|---|---|
| 3 | 0 | 37 | 39 | 11 |
| | 1 | 38 | 39 | 12 |
| | … | … | … | … |
| S2 | 24 | 62 | 50 | 19 |
| | 25 | 63 | 51 | 19 |
| | … | … | … | … |
| E2 | 93 | 99 | 65 | 26 |
| | 94 | 100 | 65 | 25 |
| | … | … | … | … |
| 4 | 155 | 160 | 82 | 35 |
| | 156 | 161 | 81 | 35 |

S2 < E2 (in this case)

**The modified branch is consisted of the points of 1-1-2 from indices 0 to 124, the points of the integer coordinate line between index 124 of 1-1-2 and index 94 of 2-1, and the points of 2-1 from indices 94 to 156.**

**Figure S4. Schematic of the modified algorithm for merging two overlapped branches.**
An example of using four topological variables (flag1, flag2, flag3, flag4) and four position indices (S1, E1, S2, E2) is shown.

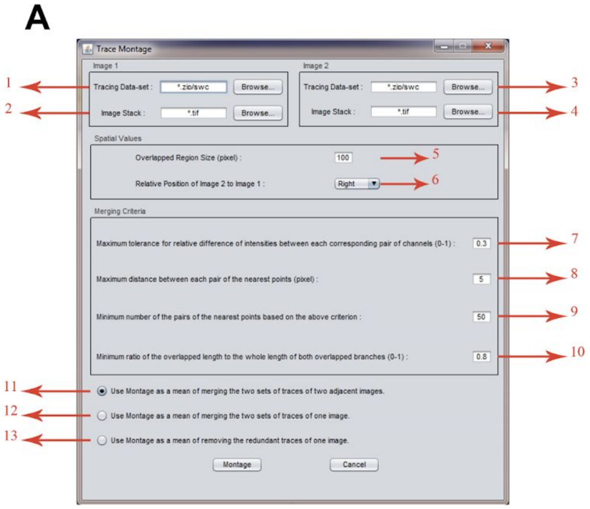


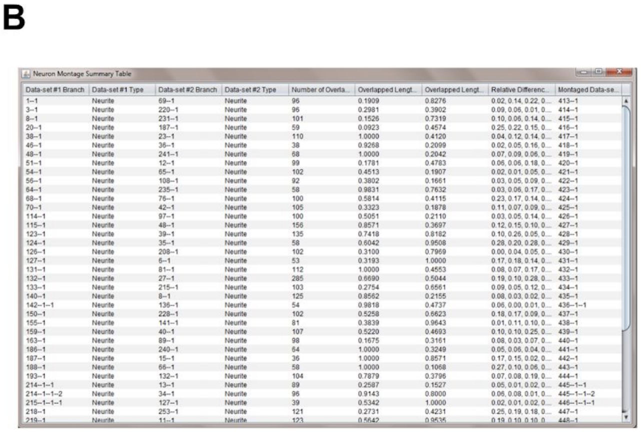


| Data-set #1 Branch | Data-set #1 Type | Data-set #2 Branch | Data-set #2 Type | Number of Overla... | Overlapped Lengt... | Overlapped Lengt... | Relative Differenc... | Montaged Data-se... |
|---|---|---|---|---|---|---|---|---|
| 1--1 | Neurite | 69--1 | Neurite | 96 | 0.1909 | 0.8276 | 0.02, 0.14, 0.22, 0.... | 413--1 |
| 3--1 | Neurite | 220--1 | Neurite | 96 | 0.2981 | 0.3902 | 0.09, 0.06, 0.01, 0.... | 414--1 |
| 8--1 | Neurite | 231--1 | Neurite | 101 | 0.1526 | 0.7319 | 0.10, 0.06, 0.14, 0.... | 415--1 |
| 20--1 | Neurite | 187--1 | Neurite | 59 | 0.0923 | 0.4574 | 0.25, 0.22, 0.15, 0.... | 416--1 |
| 38--1 | Neurite | 23--1 | Neurite | 110 | 1.0000 | 0.4120 | 0.04, 0.12, 0.14, 0.... | 417--1 |
| 46--1 | Neurite | 36--1 | Neurite | 38 | 0.9268 | 0.2099 | 0.02, 0.05, 0.16, 0.... | 418--1 |
| 48--1 | Neurite | 241--1 | Neurite | 68 | 1.0000 | 0.2042 | 0.07, 0.09, 0.06, 0.... | 419--1 |
| 51--1 | Neurite | 12--1 | Neurite | 99 | 0.1781 | 0.4783 | 0.06, 0.06, 0.18, 0.... | 420--1 |
| 54--1 | Neurite | 65--1 | Neurite | 102 | 0.4513 | 0.1907 | 0.02, 0.01, 0.05, 0.... | 421--1 |
| 56--1 | Neurite | 108--1 | Neurite | 92 | 0.3802 | 0.1661 | 0.03, 0.05, 0.09, 0.... | 422--1 |
| 64--1 | Neurite | 235--1 | Neurite | 58 | 0.9831 | 0.7632 | 0.03, 0.06, 0.17, 0.... | 423--1 |
| 68--1 | Neurite | 76--1 | Neurite | 100 | 0.5814 | 0.4115 | 0.23, 0.17, 0.14, 0.... | 424--1 |
| 70--1 | Neurite | 42--1 | Neurite | 105 | 0.3323 | 0.1878 | 0.11, 0.07, 0.09, 0.... | 425--1 |
| 114--1 | Neurite | 97--1 | Neurite | 100 | 0.5051 | 0.2110 | 0.03, 0.05, 0.14, 0.... | 426--1 |
| 115--1 | Neurite | 48--1 | Neurite | 156 | 0.8571 | 0.3697 | 0.12, 0.15, 0.10, 0.... | 427--1 |
| 123--1 | Neurite | 39--1 | Neurite | 135 | 0.7418 | 0.8182 | 0.10, 0.26, 0.05, 0.... | 428--1 |
| 124--1 | Neurite | 35--1 | Neurite | 58 | 0.6042 | 0.9508 | 0.28, 0.20, 0.28, 0.... | 429--1 |
| 126--1 | Neurite | 208--1 | Neurite | 102 | 0.3100 | 0.7969 | 0.00, 0.04, 0.05, 0.... | 430--1 |
| 127--1 | Neurite | 6--1 | Neurite | 53 | 0.3193 | 1.0000 | 0.17, 0.18, 0.14, 0.... | 431--1 |
| 131--1 | Neurite | 81--1 | Neurite | 112 | 1.0000 | 0.4553 | 0.08, 0.07, 0.17, 0.... | 432--1 |
| 132--1 | Neurite | 27--1 | Neurite | 285 | 0.6690 | 0.5044 | 0.19, 0.10, 0.28, 0.... | 433--1 |
| 133--1 | Neurite | 215--1 | Neurite | 103 | 0.2754 | 0.6561 | 0.09, 0.05, 0.12, 0.... | 434--1 |
| 140--1 | Neurite | 8--1 | Neurite | 125 | 0.8562 | 0.2155 | 0.08, 0.03, 0.02, 0.... | 435--1 |
| 142--1--1 | Neurite | 136--1 | Neurite | 54 | 0.9818 | 0.4737 | 0.06, 0.00, 0.01, 0.... | 436--1--1 |
| 150--1 | Neurite | 228--1 | Neurite | 102 | 0.5258 | 0.6623 | 0.18, 0.17, 0.09, 0.... | 437--1 |
| 155--1 | Neurite | 141--1 | Neurite | 81 | 0.3839 | 0.9643 | 0.01, 0.11, 0.10, 0.... | 438--1 |
| 159--1 | Neurite | 40--1 | Neurite | 107 | 0.5220 | 0.4693 | 0.10, 0.10, 0.25, 0.... | 439--1 |
| 163--1 | Neurite | 89--1 | Neurite | 98 | 0.1675 | 0.3161 | 0.08, 0.03, 0.07, 0.... | 440--1 |
| 186--1 | Neurite | 240--1 | Neurite | 64 | 1.0000 | 0.3249 | 0.05, 0.06, 0.04, 0.... | 441--1 |
| 187--1 | Neurite | 15--1 | Neurite | 36 | 1.0000 | 0.8571 | 0.17, 0.15, 0.02, 0.... | 442--1 |
| 188--1 | Neurite | 66--1 | Neurite | 58 | 1.0000 | 0.1068 | 0.27, 0.10, 0.06, 0.... | 443--1 |
| 193--1 | Neurite | 132--1 | Neurite | 104 | 0.7879 | 0.3796 | 0.07, 0.08, 0.19, 0.... | 444--1 |
| 214--1--1 | Neurite | 13--1 | Neurite | 89 | 0.2587 | 0.1527 | 0.05, 0.01, 0.02, 0.... | 445--1--1 |
| 214--1--1--2 | Neurite | 34--1 | Neurite | 96 | 0.9143 | 0.8000 | 0.06, 0.08, 0.01, 0.... | 445--1--1--2 |
| 215--1--1--1 | Neurite | 127--1 | Neurite | 39 | 0.5342 | 1.0000 | 0.02, 0.01, 0.02, 0.... | 446--1--1--1 |
| 218--1 | Neurite | 253--1 | Neurite | 121 | 0.2731 | 0.4231 | 0.25, 0.19, 0.18, 0.... | 447--1 |
| 219--1 | Neurite | 11--1 | Neurite | 123 | 0.5642 | 0.9535 | 0.19, 0.10, 0.10, 0 | 448--1 |

**Figure S5. *TraceMontage* user interface and quantification output.**
**(A)** The user interface, in which its different parameters (indicated by numbers) are explained in the Supplementary Methods. **(B)** The merging result table, which summarizes the montage results by showing the overlapping branches' characteristics.

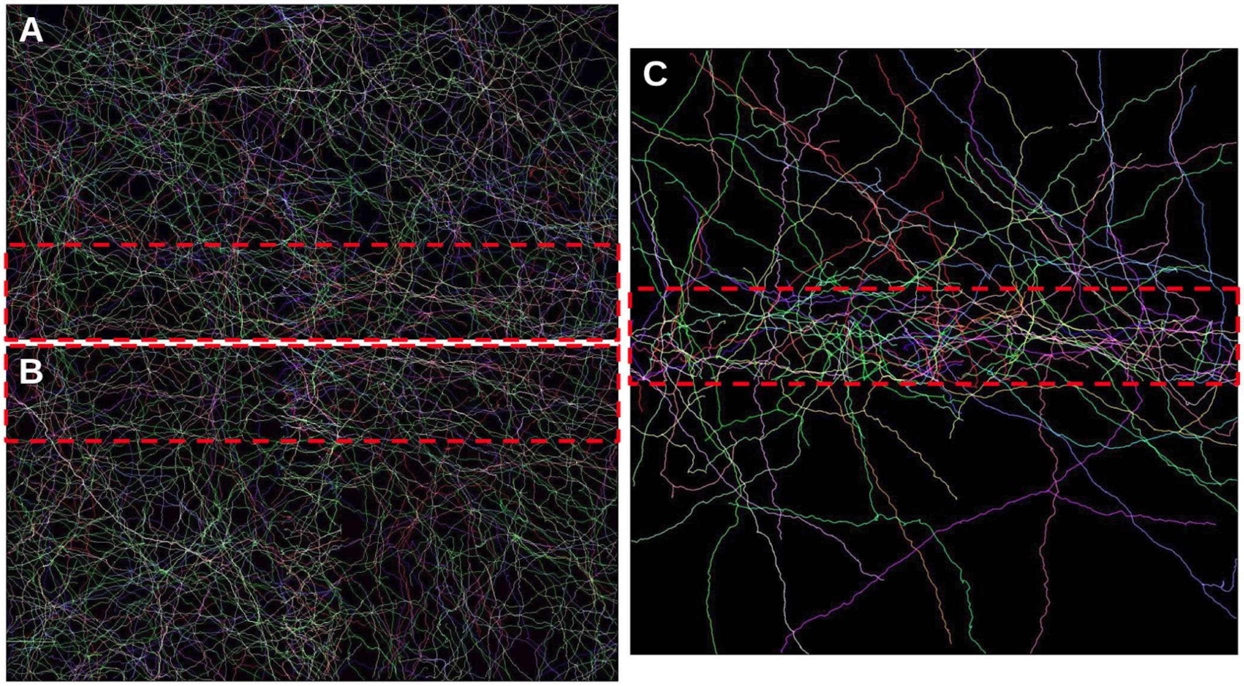


**Figure S6. A trace merging example of an image that split horizontally.**
**(A and B)** Parvalbumin-expressing basket cells in the Hippocampus of the mouse brain were imaged and split horizontally into two overlapping tiles, which were then traced independently. **(C)** *TraceMontage* was used with default settings to merge the traces. The red highlighted boxes indicate the overlapped regions of **(A and B)**.

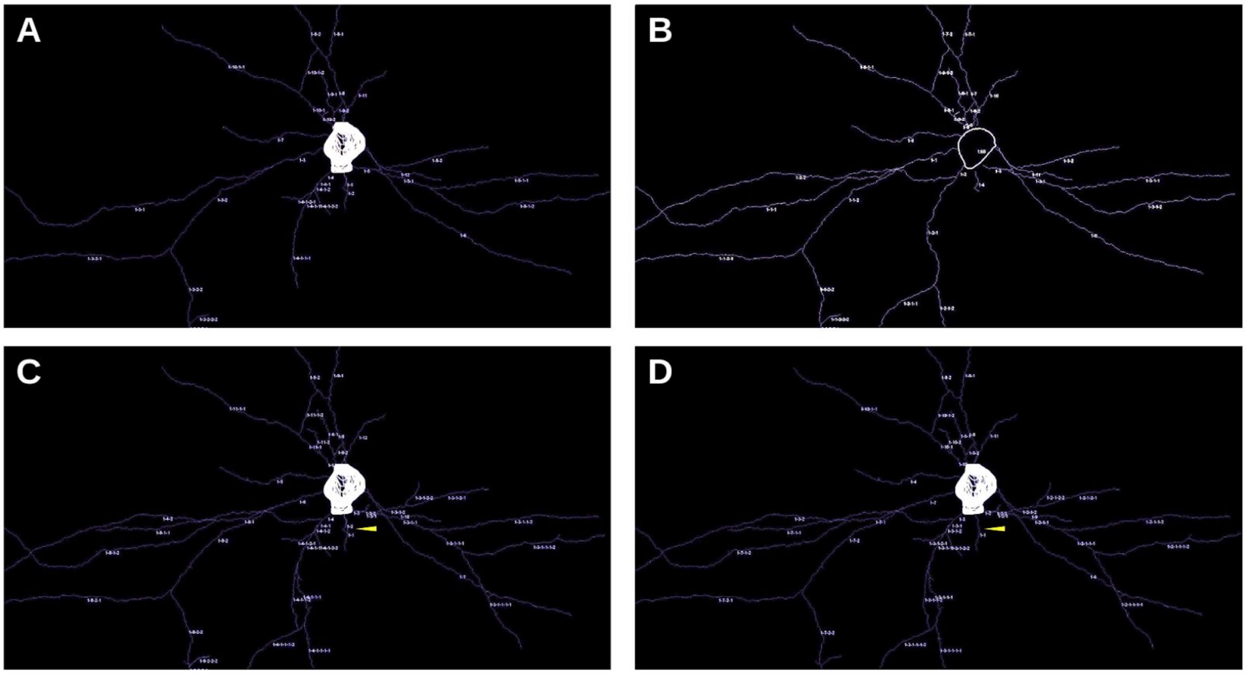

**Figure S7. An example demonstrates the merging result difference caused by selecting different input as the merging target.**
**(A and B)** Z-projection of the tracing results of the same image stack that contains a single primary visual cortex Parvalbumin-expressing basket cell reconstructed by two different tracers. **(C and D)** Z-projection of the merging results of **(A)** and **(B)** by using **(A)** or **(B)** as the target image, respectively. The default values of *TraceMontage* parameters were used in both merging processes. We can observe a small discrepancy, highlighted by yellow arrowheads, between the two montage results due to the choice of different merging targets.

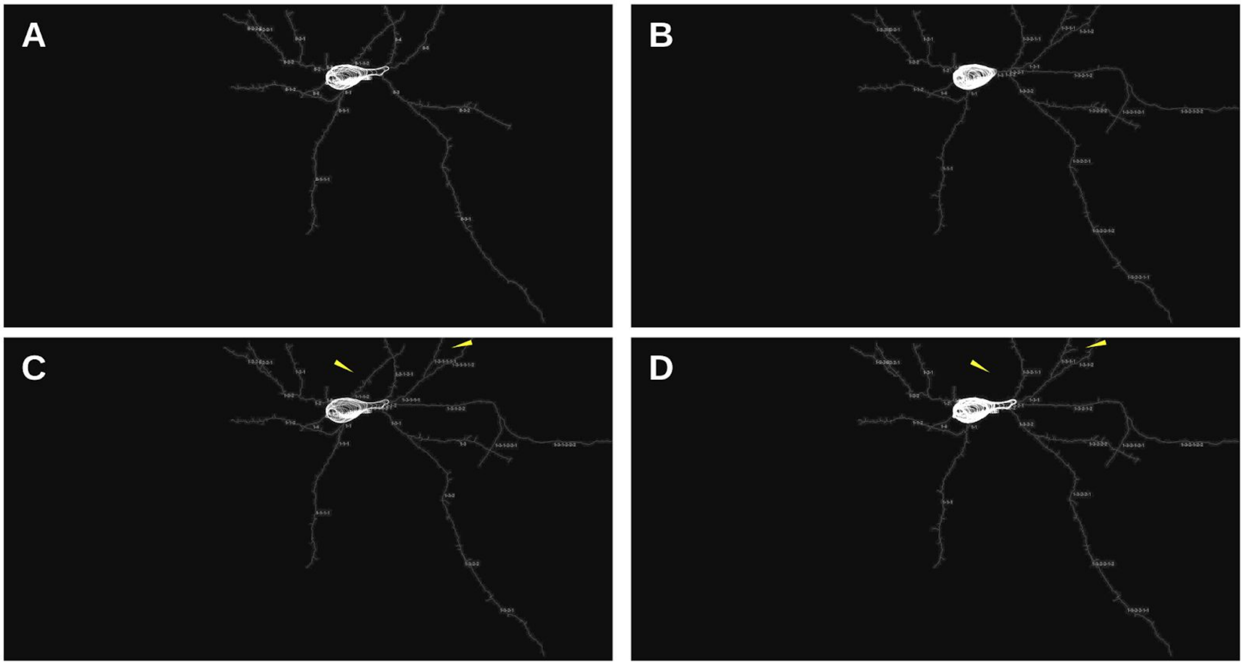


**Figure S8. Another example of merging two independent tracings of the same image by two tracers.**
**(A and B)** Z-projection of the tracing results of the same image stack that contains a single primary visual cortex Parvalbumin-expressing basket cell reconstructed by two different tracers. **(C and D)** Z-projection of the merging results of **(A)** and **(B)** by using **(A)** or **(B)** as the target image, respectively. Merging results showed differences mostly in the soma tracing (arrowheads).

### Supplemental Tables

| Column Number | Data Type | Data Value |
|---|---|---|
| **1 (N)** | Sample Number | Integer value starting from 1. |
| **2 (T)** | Structure Identifier | Standard Identifiers:<br>1. Soma<br>2. Axon<br>3. (Basal) Dendrite<br>4. (Apical) Dendrite<br>5. Fork Point<br>6. End Point<br>7. Spine |
| **3-5 (x,y,z)** | Positional Coordinates | Spatial, cartesian coordinates given in micrometers (um). |
| **6 (R)** | Radius | Half of the neurite thickness given in micrometers (um). |
| **7 (P)** | Parent Sample | The parent point's index. Parent samples should appear before any child samples. For points belonging to soma or origin points, it is -1. |
| **8 (S)** | Synapse Flag | 1 identifies a point as a synapse and 0 identifies the opposite |

**Table S1.** The *nTracer* tracing data format (modified from the SWC format) is used by *TraceMontage*. The modified format adds the eighth column in addition to the standard format to indicate the presence of a synapse.

### Supplemental Methods

The goal of neuron tracing is to provide a complete morphological reconstruction of a neuron, using a minimal and consistent model. Here, we follow the definition provided in Peng *et al.*, 2011, which defines a reconstruction of a neuron as a set of topologically connected structural components that describe the 3-D spatial coordinates of a neuron in a 3-D image. Structural components include neuron branches and other sub-structures, such as junctions and terminations, where their topological relationships are normally described through graphs (Radojević et al., 2016). In a reconstruction, graphs are often used to describe the topological relationship of all reconstruction nodes. Usually, a neuron reconstruction is described as a tree graph, which has a root node, leaf nodes, branching nodes, and other inter-nodes (Peng et al., 2011). In **Figure S1A**, the graphical representation of structural components of the input and output of *TraceMontage* is shown. It is composed of soma, subsidiary tree structure (axons, basal dendrites, apical dendrites), spines, and synapses. This structural format is the same as the structural format of output utilized in *nTracer (Roossien et al., 2019)*, which was particularly developed for the neuron tracing of Brainbow images. In the following, first, a general methodology of the montage algorithms is explained, then, an initial argument is made about their validity, and finally, the three main trace montage algorithms are described.

#### General Methodology

*Here, the general approach of the montage algorithms is explained:*

- The montage algorithms were designed to use the information provided by two input tracing datasets. The two multi-channel image stacks are solely used to calculate the colors of the branches which are used as a criterion to find the overlapped branches. As a result, the montage algorithms do not obtain any extra tracing information from those images.
- The montage algorithms use the information obtained from two input tracing datasets asymmetrically. One tracing result is set as the “target trace”, which means given a priority when tracing discrepancy is identified between the two tracing inputs. As a result, the final montage tracing result might be different when assigning different input as the “target trace”.
- The montage algorithms were designed to merge the coordinates of neurites (axons, basal dendrites, apical dendrites) that have a full binary tree structure. For somas (z-slice contours) and spines (without a tree structure), the montage algorithms solely append the coordinates of them in the final montage tracing dataset.
- There are three slightly different methods to montage the input tracing datasets: the first one assumes that two input tracing datasets belong to two adjacent images; the second one assumes that both inputs are obtained by tracing the same image; the third one simply removes the redundant traces of one input tracing dataset. The difference between the first and second methods is that the first method, after transferring the coordinates of the second input tracing dataset, considers the traces in the overlapped region of the two images to find and merge the overlapped branches. Additionally , the difference between the second and third methods is that the third method makes two copies of one input tracing dataset, and then compares them without considering the same branches to find and merge the overlapped branches.

**Derivation Arguments**

*In this section, some of the underlying logic of the montage algorithms is explained:*

**Assumption 1:** Each neuron consists of some neurites in the form of tree branches, which have a full binary tree structure. This means that each branch is either a leaf or a parent branch having exactly two child branches; e.g. the branch "1-1-2" has either no child branch or two child branches and in the latter case, the coordinates (x, y, z) of the end-point of the branch "1-1-2" is the same as the start-points of the child branches "1-1-2-1" and "1-1-2-2".

**Assumption 2:** The start point and the end point of each branch are distinct, therefore each branch does not form a loop. In addition, no two branches share the exact start and end points. As a result, the coordinates (x, y, z) of all the start and end points can uniquely determine the tree structure, including branching order.

**Lemma 1:** Based on the above two assumptions, each neuron tree structure model has exactly "n" (odd) branches (edges) with "n+1" (even) end-points (nodes). As a result, it is possible to reconstruct the whole neuron tree structure model by knowing just the relations between all edges and nodes (**Figure S2A**).

**Definition 1:** If a tracing result captures fully the tree structure of neuronal processes, we call it an "accurate" and "complete" tracing result (**Figure S2B**). A tracing result can be "accurate" but missing portions of tracing, i.e., "incomplete" (**Figure S2C**). More commonly, a tracing result can be "inaccurate" and "incomplete" (**Figure S2D**).

**Theorem 1:** If all tree branches of two input tracing results are "accurate", the montage algorithms guarantee to merge thoroughly the two input tracing datasets without missing any branches belonging to at least one of them. As a result, the final montage tracing dataset is also "accurate" and it is the union of all branches of two input tracing datasets.

**Theorem 2:** If at least one of the tree branches of two input tracing results is "inaccurate", the montage algorithms merge the maximum possible number of branches belonging to those tree branches by observing **Lemma 1**. As a result, the final montage tracing result might be "inaccurate" and it is not necessarily the union of all branches of two input tracing datasets. This might cause some branches, which exist in the input tracing results, to be missing from the montage tracing result.

**Algorithm Descriptions**

*TraceMontage* executes three main algorithmic steps obeying the principles described in the above-mentioned **Derivation Arguments** (**Figure 1**). The three main algorithms are (i) the finding algorithm (**Figure S3**), (ii) the modifying algorithm (**Figure S4**), and (iii) the merging algorithm. The code implementing each of these steps is represented in **Figure S1B**.

**The Finding Algorithm** finds the unique pairs of overlapped branches from the two input tracing results. This requires finding the overlapped branches from two tiled overlapped images (method 1), finding the overlapped branches of two tracing results obtained from the same image (method 2), or finding the redundant traces in one tracing result (method 3). In the following, the steps of the algorithm are explained (summarized in **Figure S3**):

1. Based on the relative position of the “source” image to the “target” image (assumed to be positioned to the right or the bottom) and the overlapped region size of two images, the coordinates of all branches of the “source” tracing dataset are transformed to match to the reference frame of the “target” image. For method 3, the tracing result is set as the “target” and an exact duplicate of the tracing result is set as the “source”.
2. The neurite branches which have at least one point in the overlapped region of the two images are selected from both tracing results. For method 3, all branches are selected.
3. There is a double-loop which goes through all possible pairs of selected branches of the input “target” and “source” tracing results to determine whether any given pairs remains being pre-defined as “non-overlapped”. In method 3, the same branches are excluded from pairing in this survey:
    a. The relative intensity differences between each corresponding image channel of each pair of selected branches are compared to the maximum intensity tolerance. If the intensity difference is smaller than the tolerance, the algorithm goes to step b, otherwise quits the loop.
    b. The Euclidean distance of the nearest points between the two selected branches is compared to the maximum distance tolerance. If the distance difference is smaller than the tolerance, the algorithm goes to step c, otherwise quits the loop.
    c. If the proportion of data points within either branch to the total points of both branches is greater than the minimum data proportion tolerance, the two selected branches are declared as being “overlapped”.
4. Three matching variables are defined for all pairs of “overlapped” branches (for more explanation of these three variables, please refer to the **Supplementary Program Description**):
    - “matchedOverlappedTraceBranch”
    - “matchedOverlappedTraceAnalysis”
    - “matchedOverlappedTraceCoordinate”
5. The goal of the montage algorithms is to preserve the topological features of overlapped branches. To this end, the algorithms exploit the fact that the set of two end-points of all branches belonging to a tree branch is unique, which makes us able to reconstruct the tree branches of a neuron using these endpoints. As a result, the preservation of these endpoints is important during the implementation of the algorithm. For this purpose and for each overlapped branch of the “target” tracing result, a set of all overlapped branches from the “source” tracing result is created. Among each set, the longest “source” branch that has both end-points closed, i.e., both end-points belong to other branches too, is determined as the “source” overlapped branch.
    - If none of the “source” branches have both end-points closed, the longest one is determined as the “source” overlapped branch.
6. All remaining branches of the set are removed from three main matching variables.
7. The previous two steps are repeated to select the corresponding “target” overlapped branch of each overlapped “source” branch.
8. If method 3 is used, due to the symmetric structure of the three matching variables, step 7 can be skipped.
9. Four topological variables (flags 1-4) are defined, which show the relative positioning of the two end-points of each one of the two overlapped branches, i.e., the start point of an

overlapped branch could be the end point of 0 or 1 other branch, and the end point of an overlapped branch could be the start points of 0 or 2 other branches. Also, four indices (S1, E1, S2, E2) are defined here, which indicate the index-position of the coordinates of the start points and end points of the overlapped length of the two overlapped branches in their corresponding coordinate arrays.

10. The Modifying algorithm (below) is then run, which modifies the coordinate, type, radius, synapse, and connection of the "target" overlapped branch based on the corresponding properties of the "source" overlapped branch.

**The Modifying Algorithm** modifies the coordinate, type, radius, and synapse tag of the overlapped "target" branches based on the "source" properties. It implements this by preserving the two endpoints out of four endpoints of two overlapped branches and joins the points between those two preserved endpoints. Along the modified branch and at each point with spine or connection, it records the relevant information from the corresponding overlapped branch. Letting the "target" and "source" branches be referred to as Branch1 and Branch2, respectively. And using the four topological variables (flag1, flag2, flag3, flag4) and the four indices (S1, E1, S2, E2) calculated above, the decision tree is as follows (**Figure SI4**):

- If flag1 = 1 and flag2 = 2:
  - Preserve Branch1.
- If flag1 = 1 and flag2 = 0:
  - If S2 <= E2:
    - Preserve two end-points 1 & 4
    - Join Branch1 to Branch2
    - Update Branch1
  - If S2 > E2:
    - Preserve two end-points 1 & 3
    - Reverse the order of the points of Branch2
    - Join Branch1 to Branch2
    - Update Branch1.
- If flag1 = 0 and flag2 = 2:
  - If S2 <= E2:
    - Preserve two endpoints 2 & 3
    - join Branch1 to Branch2
    - update Branch1
  - If S2 > E2:
    - Preserve two endpoints 2 & 4
    - Reverse the order of the points of Branch2
    - Join Branch1 to Branch2
    - Update Branch1
- If flag1 = 0 and flag2 = 0 and flag3 = 1 and flag4 = 2:
  - Replace Branch1 by Branch2
- If flag1 = 0 and flag2 = 0 and flag3 = 1 and flag4 = 0:
  - If S2 <= E2:
    - Preserve two endpoints 2 & 3
    - Join Branch1 to Branch2
    - Update Branch1
  - If S2 > E2:
    - Preserve two endpoints 1 & 3

        - Reverse the order of the points of Branch2
        - Join Branch1 to Branch2
        - Update Branch1
- If flag1 = 0 and flag2 = 0 and flag3 = 0 and flag4 = 2:
    - If S2 <= E2:
        - Preserve two endpoints 1 & 4
        - Join Branch1 to Branch2
        - Update Branch1
    - If S2 > E2:
        - Preserve two end points 2 & 4
        - Reverse the order of the points of Branch2
        - Join Branch1 to Branch2
        - Update Branch1
- If flag1 = 0 and flag2 = 0 and flag3 = 0 and flag4 = 0:
    - If S1 + E1 <= LengthBranch1 / 2 & S2 <= E2:
        - Preserve two endpoints 1 & 4
        - Join Branch1 to Branch2
        - Update Branch1
    - If S1 + E1 <= LengthBranch1 / 2 & S2 > E2:
        - Preserve two end points 1 & 3
        - Reverse the order of the points of Branch2
        - Join Branch1 to Branch2
        - Update Branch1
    - S1 + E1 > LengthBranch1 / 2 & S2 <= E2:
        - Preserve two end points 2 & 3
        - Join Branch1 to Branch2
        - Update Branch1
    - S1 + E1 > LengthBranch1 / 2 & S2 > E2:
        - Preserve two end points 2 & 4
        - Reverse the order of the points of Branch2
        - Join Branch1 to Branch2
        - Update Branch1

**The Merging Algorithm** generates the output tracing results following the input tracing data format by removing, reordering, and renaming the branches (neurite, soma, spine) and modifies the synapse variable. It guarantees the following:

1. Those neurons (consisted of neurites, somas, and spines with all properties) without any overlapped branches from the input tracing results are saved in the montage variables after renaming their branches. The tag property of these neurons are modified to show that the neurons initially belong to which input tracing result.
2. For each pair of the overlapped branches, the pair of tree-branches which they belong and their corresponding pair of neurons are determined with all of their branches (neurite, soma, spine). This new set of branches is going to be organized into a new single neuron after removing, reordering, and renaming of the branches.
3. The tag property of the new neuron is modified to show that the new neuron is the result of merging two of the input traces.
4. Soma contours are added to the montage variables after renaming. Priority is given to the data obtained from the “target” tracing result.

5. All the non-overlapped neurites are added to the montage variables after renaming.
6. Overlapped branches are reorganized into a new set. After modification, the maximum possible number of the neurite branches is selected to build a new tree following rules established in **Lemma 1**. The new tree branches will be renamed with all properties preserved.
7. The spines are added to the corresponding modified branches in the montage result.
8. The synapse tags are modified similar to the spines.

**Notes on output:** The following statistical measurements are saved in a Microsoft Excel file:
1. Ratios of merged branches to all branches in both tracing datasets.
2. The ratio of merged branches with similar types to all merged branches.
3. The means and standard errors of the number of overlapped points of the pairs of merged branches of both inputs.
4. The means and standard errors of the ratio of the overlapped length to the whole length of the merged branches of both inputs.
5. The means and standard errors of the relative difference of intensities among each image channel of all overlapped branches.
